# Tracking the stochastic growth of bacterial populations in microfluidic droplets

**DOI:** 10.1101/2021.01.08.425714

**Authors:** Daniel Taylor, Nia Verdon, Peter Lomax, Rosalind J. Allen, Simon Titmuss

**Author notes:** DT and NV made equal contributions to this work. RJA and ST made equal contributions to this work.

## Abstract

Bacterial growth in microfluidic droplets is relevant in biotechnology, in microbial ecology, and in understanding stochastic population dynamics in small populations. However, it has proved challenging to automate measurement of absolute bacterial numbers within droplets, forcing the use of proxy measures for population size. Here we present a microfluidic device and imaging protocol that allows high-resolution imaging of thousands of droplets, such that individual bacteria stay in the focal plane and can be counted automatically. Using this approach, we track the stochastic growth of hundreds of replicate *Escherichia coli* populations within droplets. We find that, for early times, the statistics of the growth trajectories obey the predictions of the Bellman-Harris model, in which there is no inheritance of division time. Our approach should allow further testing of models for stochastic growth dynamics, as well as contributing to broader applications of droplet-based bacterial culture.

## Introduction

Droplet microfluidics has a plethora of well-established applications in biology, ranging from highly sensitive detection methods (e.g. droplet-based polymerase chain reaction [1]) to high-throughput optimisation of chemical or enzymatic activities [2–4]. The ability to cultivate biological cells in droplets opens up further applications [5, 6], such as high-throughput screening for gene expression [7, 8] or metabolite production [9], screening of antibodies [10] or of environmental isolates for drug discovery [11], understanding microbial interactions [12–16], investigation of responses to drugs such as antibiotics [17–21] and the detailed study of stochastic growth dynamics and evolution [22–25].

Many challenges need to be overcome when growing biological cells within droplets in a microfluidic device. Encapsulation of cells in droplets is inherently a random process, that can be well described by Poisson statistics [26]. Moreover the cells need to be maintained at a fixed temperature and supplied with appropriate nutrients including oxygen. One may wish to change the environmental conditions during the experiment (e.g. by infusing a drug) or across the device, and one may need to sort the droplets at the end of the experiment. All of these challenges have been addressed through appropriate microfluidic device design [6, 8, 9, 17, 25, 27–31].

Less attention has been paid, however, to the problem of how to directly track population growth within microfluidic droplets. Most microfluidic droplet methodologies do not produce sufficiently high-resolution images to allow individual cells to be counted as they proliferate. Therefore, growth is often tracked using the total droplet fluorescence as a proxy for population size [6, 11, 17, 20, 22, 24, 32, 33] – assuming that the cells are fluorescent, or can be made so. This approach is especially widely used for bacteria, whose cells (of size ∼ 1-5*µ*m) are much smaller than those of eukaryotes (e.g. yeast or mammalian cells). Alternatively, light scattering from individual droplets has been used as a proxy measure for bacterial growth [29], that is analogous to the optical density measurements used for bulk cultures. However, the fluorescence intensity of individual cells can vary significantly across a population [34], and can also change with growth state (e.g. entry to stationary phase [35]) – while light scattering is also sensitive to factors such as cell size and shape, which can be affected by drug exposure or other changes in environment [36]. Thus, it is hard to reliably link these proxies to cell number. This is important when studying stochastic population dynamics, where cell number fluctuations are the key quantity of interest [22, 37]. While some studies have measured cell numbers directly in microfluidic droplets [12, 14, 15, 19, 23], this has typically been done by manual counting, which is labour-intensive, especially for bacteria. Recently, an innovative microfluidic hemocytometer approach has also been used to count individual bacteria as well as human cells in droplets [38]; however this method is not suitable for tracking growth dynamics within individual droplets.

Bacterial growth and division are intrinsically stochastic processes. This individual cell stochasticity translates into stochastic population growth, with implications e.g. for survival of antibiotic treatment [39–42]. From a theoretical point of view, there has long been interest in growth of populations in which individual cell lifetimes are stochastic [43– 46], but it is challenging to distinguish alternative models with existing experimental data [22]. Tracking individual bacteria growing in multiple replicate droplets can improve our ability to link growth stochasticity at the individual cell level to the population level, and to test different theoretical models for stochastic population dynamics.

Here, we present a microfluidic device and image analysis protocol that allows automated, high-throughput *counting* of individual fluorescent bacteria in multiple replicate microfluidic droplets. This allows extraction of growth trajectories within each droplet, which show significant droplet-to-droplet variation for the bacterium *Escherichia coli* in glucose minimal medium. Our device relies on the vertical squashing of droplets within the microfluidic reservoir where they are imaged, into discs of thickness 11.5*µ*m (Figure 1(aiii)), such that bacteria remain in the focal plane as they proliferate. This is combined with an image analysis workflow that includes droplet detection, tracking, thresholding and spot counting. Analysing our results for stochastic growth of hundreds of *E. coli* populations on glucose minimal media, we find good agreement at early times with the predictions of the Bellman-Harris model [46], in which there is no inheritance of division time. We infer the mean and variance of individual cell division times from the stochastic population growth trajectories, which had proved impossible in previous work [22]. Our approach should allow further testing of models for stochastic growth dynamics, as well as being contributing to applications such as antibiotic discovery [11].

**FIG. 1:**
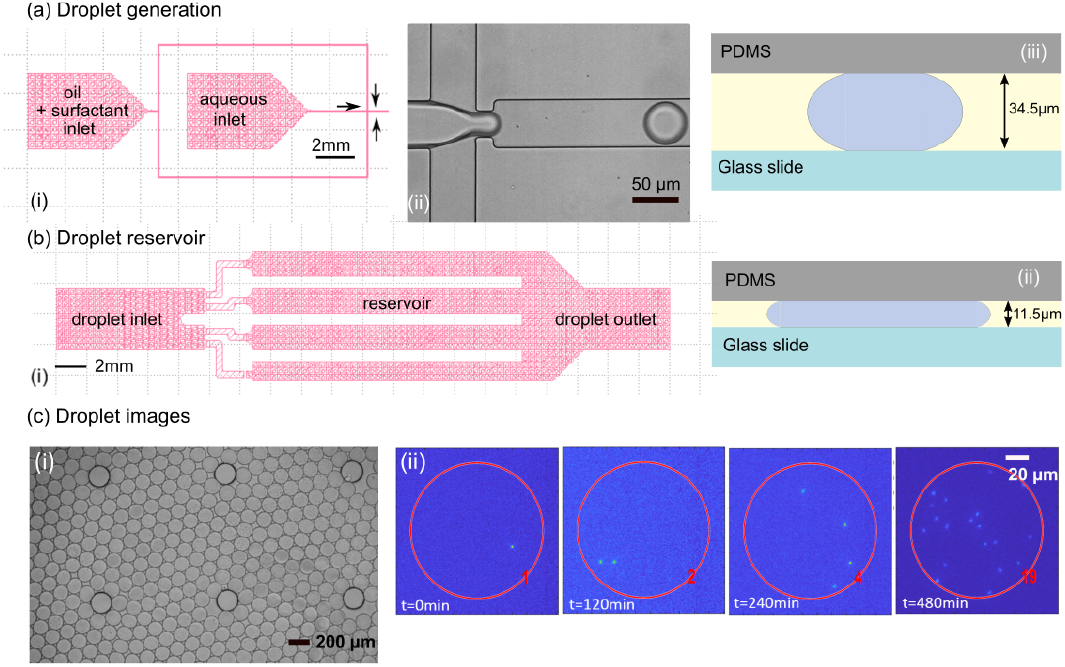
Device design. Droplets are rapidly generated using a microfluidic device with flow-focusing geometry (a). The oil and aqueous phases are pumped into the device, shown in (ai), using syringe pumps. (aii) shows a droplet as it is generated; (aiii) shows a scale drawing of a droplet of volume 85 pL in the generator. Droplets are then transferred to a reservoir (b), which has a reduced channel height, squashing the droplets (bii) to allow microscopic imaging within the field of focus. The diagram in (bii) is to scale, such that a droplet of volume 85 pL would have diameter 100*µ*m in the reservoir. (c) shows droplets in the brightfield (ci) and fluorescence (cii) channels. (cii) shows multiple fluorescence images of the same droplet at different times during the experiment. The red circle represents the droplet outline that has been detected from the brightfield image. The number of bacteria detected in this droplet by our automated counting algorithm is indicated in red; this number increases in time as the bacteria proliferate within the droplet.

### Device development and validation

#### A microfluidic device for imaging growth of individual bacteria in droplets

Our microfluidic device generates a low polydispersity emulsion of aqueous phase droplets in a fluorocarbon continuous phase and stores them in a reservoir, allowing observation of encapsulated bacterial communities using brightfield and fluorescence microscopy. The device design is shown in Figure 1. The device is formed from polydimethylsiloxane (PDMS) bonded to a glass slide, and consists of two parts: a droplet generator and a droplet reservoir.

The droplet generator uses a flow-focusing junction with channels of width 50*µ*m that come together in a 20*µ*m constriction in which the aqueous phase (bacterial suspension) in the central channel is squeezed between the two outer oil phase flows [47] (Figure 1a(i),(ii)). This is followed by a serpentine channel which allows mixing within the droplets (not shown in Figure 1). The droplet generator is 34.5*µ*m in height; for the flow rates used in our work, this leads to production of droplets of mean volume 120pL, with a standard deviation of 14pL. We note that when unconfined, the radius of a 120pL droplet is greater than the channel depth of 34.5*µ*m, so that the droplets are compressed when generated, as shown in Figure 1a(iii). After generation, droplets are transferred from the generator to the reservoir via a piece of tubing of length 2-2.5cm (Tygon S3 E-3603, inner/outer diameter 0.8mm/2.4mm).

The droplet reservoir (Figure 1b(i)) receives an input flow of droplets from the generator, and stores them in an array of 4 parallel channels. The parallel channel design, as well as the inlet geometry and teardrop-shaped support pillars, aims to disperse droplets laterally, allowing efficient filling of the reservoir (Figure 1c). Droplets can freely exit the reservoir via a large outlet, preventing jamming. Once the inflow of droplets stops, droplet movement within the reservoir is arrested almost completely, which proves helpful for droplet tracking. Importantly, the small reservoir height (11.5*µ*m) ensures that droplets are significantly squashed in the vertical direction (Figure 1b(ii)); this prevents bacteria from moving out of the microscope focal plane, allowing them to be accurately counted. We have verified that the bacterial count remains stable when we scan the reservoir over a distance of ∼ 12*µ*m in the vertical direction (Figure S1). We aim for a loose packing of the disk-shaped droplets within the reservoir, to prevent shape deformation or adhesion between droplets. Typically, in our experiments the reservoir contains approximately 1000 droplets (Figure 1c(i)). The entire device (droplet generator, tubing and reservoir) is maintained at a fixed temperature of 37°C by being submerged under water in a custom 3D-printed microscope mount coupled to an external temperature-controlled water bath. As well as maintaining temperature, submerging the device in water suppresses evaporation from the aqueous droplets [48].

In this work, droplets are created using as the aqueous phase a dilute suspension of *Escherichia coli*, strain RJA002, which constitutively expresses yellow fluorescent protein (YFP) and therefore fluoresces yellow (see Methods). The oil phase consists of oxygen-permeable, biologically inert FC40, containing the surfactant Pico-Surf at 2.5%*w/w* [59].

#### Imaging bacterial growth in droplets

To analyse bacterial growth within the droplets, the entire reservoir is imaged every 6 minutes over a period of 8 hours. Each image is recorded both using brightfield illumination and in the YFP fluorescence channel (Figure 1(c)). To count individual bacteria, high magnification images are required, typically 20× and above, implying that multiple fields of view are needed to cover the entire reservoir. The reservoir is therefore imaged in a grid pattern at 20× magnification, and the fields of view are then stitched together. Each composite image consists of 18×14 fields of view, with each field of view containing up to 25 droplets. In an 8h experimental run, each grid position is imaged 80 times.

Once the experimental run is complete, all the images are checked manually. Fields of view that are out-of-focus, do not contain any droplets (e.g. because the region imaged lies between reservoir channels), contain distorted or non-circular droplets, or are contaminated with debris, are replaced with blank image stacks over the entire time course of the experiment.

#### Image processing: droplet identification and tracking

After stitching together and checking the fields of view, we obtain two composite image stacks, one for brightfield and the other for fluorescence, containing all fields of view over all time steps. Next, droplets are detected in the brightfield image using a circular Hough transform algorithm (see Methods), which fits circular boundaries to the droplets and outputs their centre positions and radii (Figure 2(a)). The circles detected by the algorithm are considered to be droplets if their radii lie between certain bounds; this allows for the exclusion of rare large droplets and circular features in the microfluidic device structure, such as reservoir supports. Droplet detection is performed for every timestep, leading to a dataset containing the position and radius of every droplet detected at each timepoint.

**FIG. 2:**
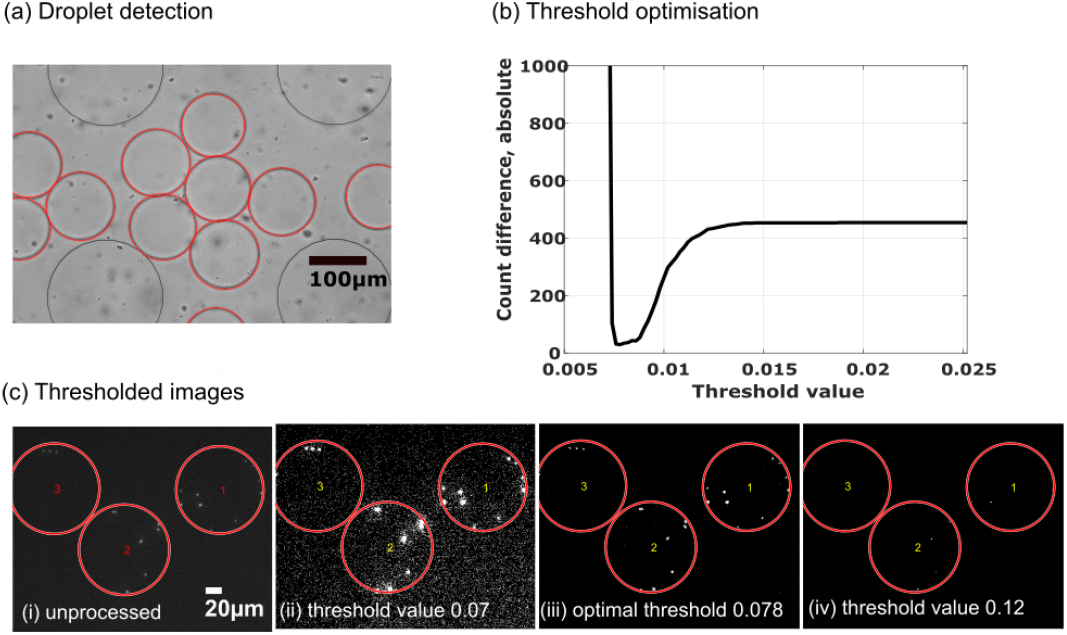
Image processing. (a) Droplet detection using the circular Hough transform algorithm; the circles that are identified as droplets are shown in red. (b) Optimisation of the binarisation threshold for detection of individual bacteria in fluorescence images. The absolute difference between the total number of bacteria recorded by the algorithm and a manual count is plotted as a function of the threshold pixel intensity value. Here images were taken at 120 mins and comprised 118 droplets with 475 bacteria recorded in the manual count. (c) Binarised fluorescence images (with overlaid droplet boundaries) obtained using a range of threshold values. If the threshold value is too low, spurious small objects are detected; if it is too high bacteria are missed.

To measure bacterial population growth within a given droplet, we track droplets from one time step to the next using an adapted particle tracking algorithm [49]. This works well because very few droplets move by more than their own radius during the 6 minute time interval between images.

Upon completion of both the droplet boundary detection algorithm and the droplet tracking algorithm, a data array is output. This array contains the centre position and radius of every droplet across all time steps. In the array, each droplet is also assigned an identification number that allows it to be followed in time, from one frame to the next.

#### Image processing: counting individual bacteria

To count individual bacteria within a droplet, the fluorescence images are binarised (see Methods). The binarisation threshold is optimised on a subset of images (4 fields of view, at a timepoint midway through the experiment) to minimize the absolute difference between a manually conducted count, and our automated counting algorithm (see Methods and Figure 2(b) and (c)). Once the optimal threshold value has been determined, all of the images are binarised using this threshold and then passed to the automated bacterial counting algorithm. This counts the number of distinct fluorescent objects within each droplet boundary, for each image. Using the droplet identification numbers from the tracking algorithm, it is then possible to track changes in bacterial numbers in specific droplets, over the duration of a complete experimental run (see, e.g. Figure 1(cii)).

The bacterial counting algorithm was found to be accurate to within ∼ 4.7%, when compared to manual counts for a sample of 68 droplets, containing on average 7 bacteria (droplets with zero bacteria were excluded from this analysis). To analyse in detail the source of counting errors, we also analysed a larger dataset consisting of 118 droplets, with a total of 475 bacteria, imaged after 120min of growth. Here we found 17 droplets for which the automated bacterial count differed from the manual count. Of these, the automated algorithm counted too few bacteria in the majority of cases (14/17). The magnitude of the count error did not correlate with either the number of bacteria in the droplets (counted manually), or the total droplet fluorescence (Figure S2(a)), suggesting that there is not a simple link between bacterial density and count error, for early-time trajectories. We also looked in detail at the images for these 17 droplets, to characterise the source of error (Figure S2(c) and (d)). In 6/17 cases, errors were caused by clumping of bacteria, causing multiple bacteria to be counted as one unit. In 7/17 cases we observed that two daughter cells had just separated but are counted by the algorithm as one unit; this is a biological process rather than an error as such. The few remaining errors were caused by random pixels being counted as bacteria (1/17), an incorrect droplet boundary position (1/17), or fluorescent dust particles (2/17).

#### Device validation

To confirm that the bacterial population in each of the droplets behaves independently, we measured the spatial correlation between droplet populations. We construct a correlation metric *S*(*r*_*ij*_), for a given timepoint (here 480min), according to the following procedure. For each pair of droplets (*i, j*) we measure the centre-to-centre distance (*r*_*ij*_), and the absolute difference in population size (|*P*_*i*_ −*P*_*j*_|), between the droplets. Pairs of droplets (*i, j*) are sorted into bins on the basis of their centre-to-centre distance *r*_*ij*_, and we compute the mean difference in population size for the pairs of droplets in each bin (i.e. the sum of |*P*_*i*_ −*P*_*j*_| values divided by the number of droplet pairs in the bin). This produces a spatial correlation function for the absolute difference in population size between pairs of droplets. We detected no significant spatial correlations in population size between the droplets (Figure 3(a)).

**FIG. 3:**
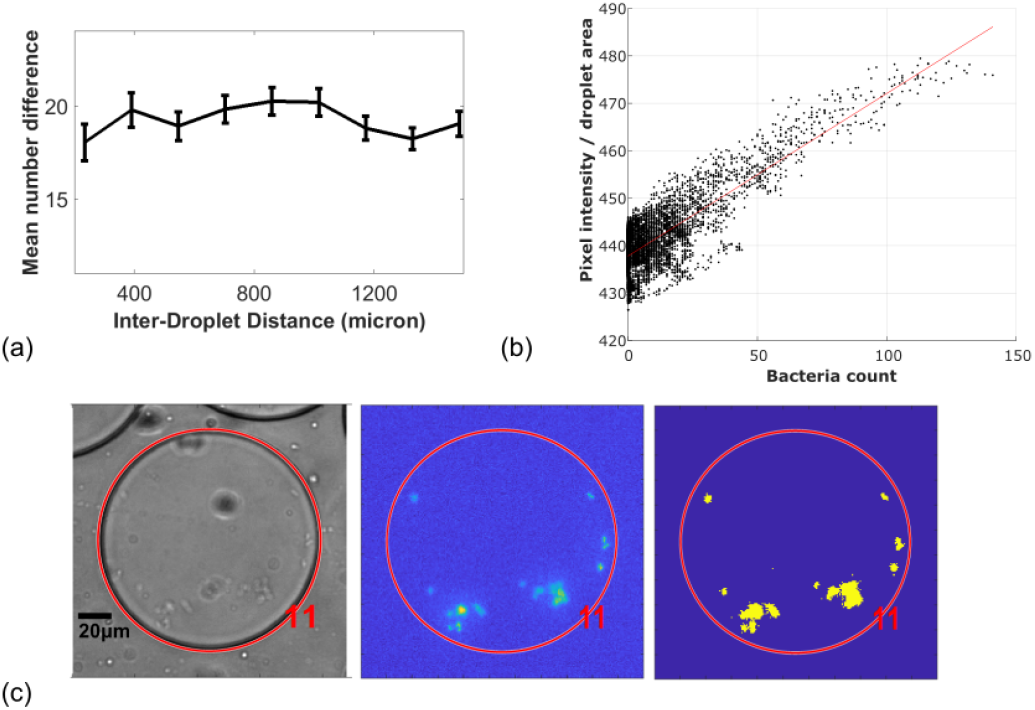
Method validation. (a): Absence of spatial correlation. Mean absolute difference in bacterial counts between pairs of droplets, plotted as a function of inter-droplet separation (i.e. mean value of |*P*_*i*_ *–P*_*j*_| for droplet pairs whose spatial separation lies within a bin assigned on the basis of the inter-droplet distance *r*_*ij*_. Data corresponds to a single timepoint at 480 min. The error bars show 95% confidence intervals for the mean (1.96*×*SEM). (b): Comparing bacterial counts measured using our algorithm to total droplet fluorescence values. The data points represent 118 droplets tracked over the entire experimental run (therefore different data points can represent different times). To evaluate total fluorescence, pixel values are summed over an entire droplet and the sum is normalised by the droplet area, to control for background fluorescence, which scales with area. Total fluorescence correlates strongly with bacterial count: Pearson correlation coefficient = 0.84 (P*<*0.001). The best fit line is *y* = 0.34*x* + 438 (*R*^2^ = 0.66). (c): Representative images showing a droplet in which bacteria have clumped: phase contrast image (left), fluorescence image (middle) and binarised fluorescence image (right). The droplet counting algorithm gives an incorrect result of 11 bacteria.

To verify that our device provides adequate nutrients and oxygen for bacterial growth, we compared the total rate of bacterial growth within the device with that measured in a bulk culture. Summing the total bacterial numbers across 188 droplets in our device, measured over a run of length 150 minutes, in M9 glucose minimal media (see Methods), we observed exponential growth with a doubling time of 89 ± 1 minutes. This is comparable to the doubling time of 84 ± 2 minutes for the same strain and media in bulk culture [50].

#### Comparing bacterial counting to total droplet fluorescence

Previous work has monitored bacterial growth in droplets using total droplet fluorescence as a proxy for bacterial number [6, 11, 17, 20, 22, 24, 32, 33]. Therefore we compared bacterial counts for individual droplets, obtained using our counting algorithm, with the integrated total fluorescence across the droplet (Figure 3(b); data from all time points are included on the plot; total fluorescence is normalised by droplet area). We observe correlation between bacterial number and integrated droplet fluorescence (Pearson correlation coefficient = 0.84, P*<*0.001), particularly for small bacterial numbers (corresponding typically to early times during the experiment), although there is significant noise in the fluorescence values for a given bacterial count. This could arise from heterogeneity in fluorescence intensity between bacteria [34]. For larger bacterial numbers, there is some indication that the integrated fluorescence plateaus, such that it no longer correlates with bacterial count. This may correspond to the late-stage growth regime, when the fluorescence intensity of individual bacteria may start to decrease [35]. This effect can be seen more clearly when comparing trajectories of total fluorescence and bacterial count for individual droplets (Figure S3).

#### Encapsulation of bacteria follows a Poisson distribution

From similar microfluidics experiments, we expect the loading of bacteria into droplets to conform to the theoretically predicted Poisson distribution [26]. To test this, we analysed the distribution of the numbers of bacteria in the droplets for 375 droplets loaded with fluorescently labelled *E. coli* bacteria in M9 minimal medium at the start of our monitoring period. At the loading density used, 179 of the droplets were empty while 196 contained bacteria.

Figure 4(a) shows the measured distribution of bacterial numbers in the droplets for our first timepoint. The blue region shows the best fit to a Poisson distribution (with 95% confidence interval), while the red region shows that a much better fit is obtained to a computational model in which we simulate exponential growth of bacteria that are initially Poisson distributed (see Methods). The fact that our data is well fitted by a model that includes Poisson loading and growth reflects the fact that, because droplets have to transfer between the generator and the reservoir, there is a time delay of ∼95 mins (approximately 1 doubling time) between droplet creation and the start of our imaging. During this time, bacteria can proliferate within the droplets.

**FIG. 4:**
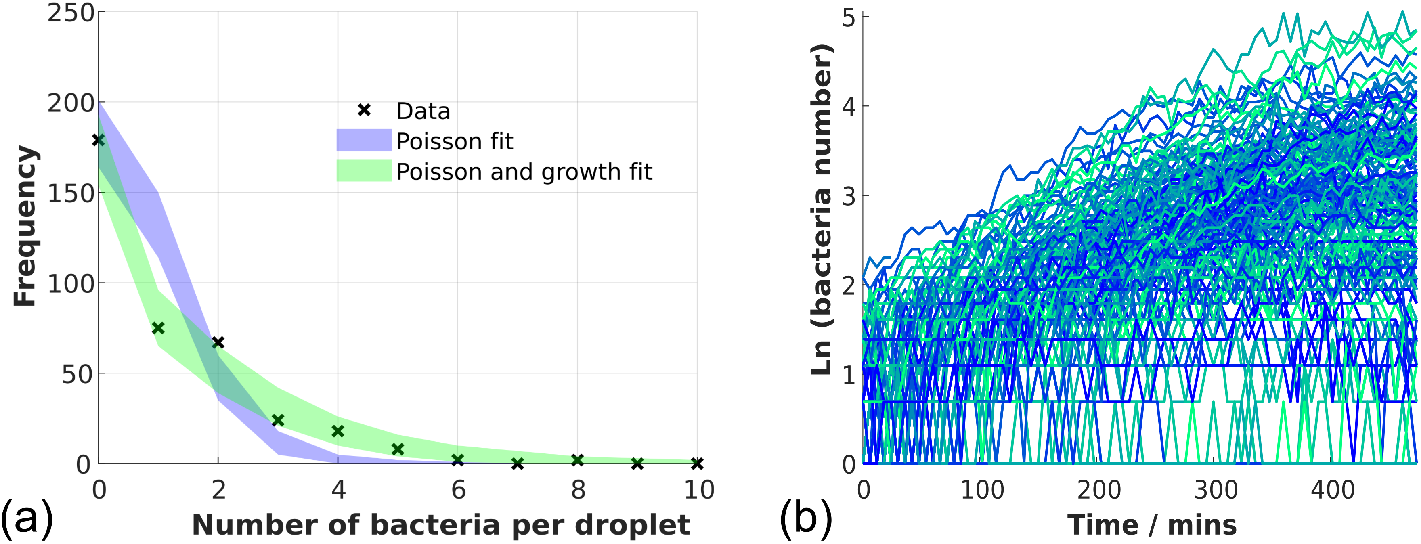
Stochastic encapsulation and growth of *E. coli* cells in droplets. (a): Distribution of droplet occupation numbers at the first imaged timepoint. The blue area shows a fit to a Poisson distribution (the shaded area shows 95% confidence interval) while the green area shows the fit to a model where initially Poisson-distributed bacteria proliferate exponentially (see Methods for more details). (b): Trajectories of logarithm of bacterial count for individual droplets; extensive variability in growth is observed.

#### Counting errors lead to transient drops in growth trajectories

To investigate stochastic bacterial growth in individual microfluidic droplets, we tracked over time the growth of the 196 *E. coli* populations contained in our 375 droplets. Figure 4 (b) shows that, although growth within the droplets is broadly exponential, growth trajectories are highly variable between droplets.

Occasionally, apparent decreases in bacterial number within individual droplets can be seen. Since spontaneous bacterial lysis is unlikely under these conditions, these are most likely caused by counting errors. To check the origin of the transient drops in the bacterial count trajectories, we analysed in detail 3 droplet trajectories that showed such drops (Figure S4). Comparing the microscopy images of these droplets before and after the transient drops in bacterial count (Figures S5, S6 and S7), we found that indeed the drops were caused by errors. In two cases (Figures S5 and S6) the error occurred because several bacteria happened to be close together and were counted as a single unit. In the third case (Figure S7), a droplet was distorted by a nearby pillar in the reservoir, causing the droplet boundary-finding algorithm to misdraw the boundary, such that a bacterium was missed in the count. In the same trajectory, two bacteria that were colocalised in space were also counted as a single unit (Figure S7).

#### Bacteria tend to clump at late times

At late times in our experiments, we observe more frequent and larger drops in the bacterial count (Figure S4). Checking our database of microscopy images, we observe that when the bacterial density in the droplets becomes high, the bacteria can form clumps (Figure 3(c); see also Figure S6); indeed a few small clumps are observed from ∼ 60 min onwards. This leads our counting algorithm to produce incorrect results – although we note that it is generally not possible to count the individual bacteria in these clumps manually either. Clumping of *E. coli* bacteria in microfluidic droplets has also been observed by other authors [21]. This issue might be avoided in future work by the use of non-clumping bacterial strains (see Discussion). From a practical point of view, the appearance of large apparent fluctuations in the bacterial count might provide a useful sign that the accuracy of the automated count can no longer be relied on, due to clumping. The beauty of our approach is that the images of the droplet corresponding to that particular trajectory can be examined to confirm that a biologically phenomenon, such as clumping is occurring. Being able to assay for clumping in this way could prove interesting, for example, in the study of biofilm initiation.

### Stochastic dynamics of bacterial growth

Our system allows us to probe in detail the statistical properties of stochastic bacterial growth trajectories in small populations. In principle, such analysis can be used to test different models for the stochastic process of single-cell growth and division, and to extract information on single-cell growth and division parameters. In recent work, Barizien *et al*. [22] analysed a dataset of growth trajectories for *E. coli* populations in microfluidic droplets, in which the readout was total droplet fluorescence (*i*.*e*. absolute bacterial counts were not available). Barizien *et al*. showed that their data was consistent with the Bellman-Harris (BH) model [46]. The BH model considers stochastic population growth, starting from exactly 1 bacterium, and assumes that the division times of all bacteria in the population are independent and chosen from a fixed probability distribution (which Barizien et al. took to be a Gaussian with mean *τ* and standard deviation *σ* [22]). In Barizien et al.’s work, three sources of stochasticity contributed to the variability in population growth dynamics: intrinsic stochasticity from the single-cell growth and division process, stochasticity in initial population size from the Poisson loading of the droplets, and changes in the overall growth rate during the experiment [22]. The latter two effects tended to obscure the signal from the intrinsic single-cell stochasticity. Our methodology provides a way to overcome this problem. Because we are able to count individual bacteria, we can group droplets in our dataset according to their initial bacterial occupancy. Analysing each group separately removes the effect of the stochastic droplet loading. Moreover, we do not observe changes in growth rate during the early stages of our experiments (perhaps because bacteria acclimatize to the device during the wait time between droplet generation and arrival in the reservoir). Applying the analysis pioneered by Barizien et al. to our dataset, we are able to infer values for the single-cell growth parameters: the mean and standard deviation of the division time *τ* and *σ*.

#### Growth dynamics are consistent with the Bellman-Harris model

The BH model makes predictions for the statistics of the number of bacteria *N* (*t*) in a population at time *t*, starting from a single bacterium at time zero, and assuming that bacteria divide independently with division times that are chosen from a fixed distribution [46]. The BH model predicts that both the mean and the standard deviation of *N* (*t*) grow exponentially with the same exponent *α*, i.e. they tend to *n*_1_*e*^*αt*^ and *n*_2_*e*^*αt*^ at long times. Hence the coefficient of variation *CV*_1_, the standard deviation of the population size *N* (*t*) divided by the mean, tends to a constant value *n*_2_*/n*_1_. The values of the constants *α, n*_1_ and *n*_2_ depend on the parameters of the underlying distribution of division times. Barizien et al. extended the work of Bellman and Harris to show that for a population initiated with *k* bacteria, the coefficient of variation *CV*_*k*_ still tends to a constant, but the value of the constant now depends on *k* [22]:

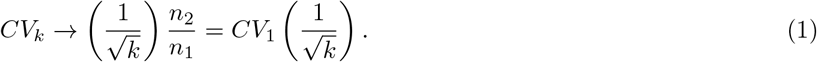

Furthermore, for the case where the initial bacterial numbers are Poisson-distributed, the coefficient of variation of the population size *CV*_*λ*_ (computed over non-empty droplets) obeys the relation [22]:

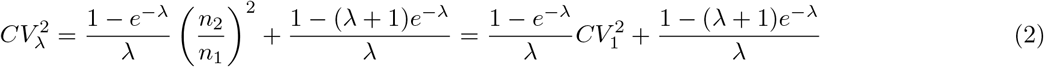

where *λ* is the parameter of the Poisson distribution, i.e. the mean initial bacterial number (computed over all droplets including empty ones).

We analysed 188 non-empty droplet trajectories for *E. coli* bacteria in M9 minimal medium supplemented with glucose (see Methods). Our complete dataset consisted of 375 droplets, of which 184 were empty (3 trajectories were manually disgarded due to the presence of imaging artefacts, e.g. dust). Of the 188 non-empty droplet trajectories, 68 initiated with 1 bacterium, 66 with 2 bacteria, 23 with 3 bacteria, 19 with 4 bacteria, and 8 with 5 bacteria. We therefore divided the trajectories into five groups according to their initial population size (omitting the 4 trajectories that initiated with more than 5 bacteria). For each group of trajectories, we computed the mean and standard deviation of the population size, and the coefficient of variation, as functions of time. Figure 5(a) and (b) show that the mean and standard deviation of the population size grow exponentially in time, regardless of initial population size, as predicted by the BH model. Figure 5(c) shows that, at early times (up to ∼ 150min), the coefficient of variation *CV*_*k*_ is indeed constant in time, as predicted. At later times, the coefficient of variation tends to increase slightly, perhaps suggesting a deviation from true exponential growth. The BH model makes the specific prediction that the coefficient of variation *CV*_*k*_ should scale with the inverse square root of *k* (Eq. (1)). Figure 5(d) tests this directly, by plotting the ratio *CV*_*k*_*/CV*_1_ (averaged over the first 150min) versus 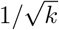. Indeed this data collapses rather well onto the predicted 1:1 straight line. We can also test the prediction of Eq. (2) for the coefficient of variation of the entire set of 188 non-empty droplets, *CV*_*λ*_. In our dataset, the mean initial bacterial number (including empty droplets) is *λ* = 1.11. Hence, from Eq. (2) we expect that 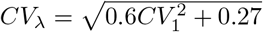. Evaluating the coefficient of variation across all the 188 non-empty droplets in our dataset, averaged over the first 150min (bold black data in Figure 5(c)), we obtain 0.681 ±0.008. This is in good agreement with the value of 0.70 ±0.05 that we obtain from right-hand side of Eq. (2), evaluated for our dataset starting with one bacterium per droplet (for which *CV*_1_ = 0.60 ±0.05, averaged over the first 150min).

**FIG. 5:**
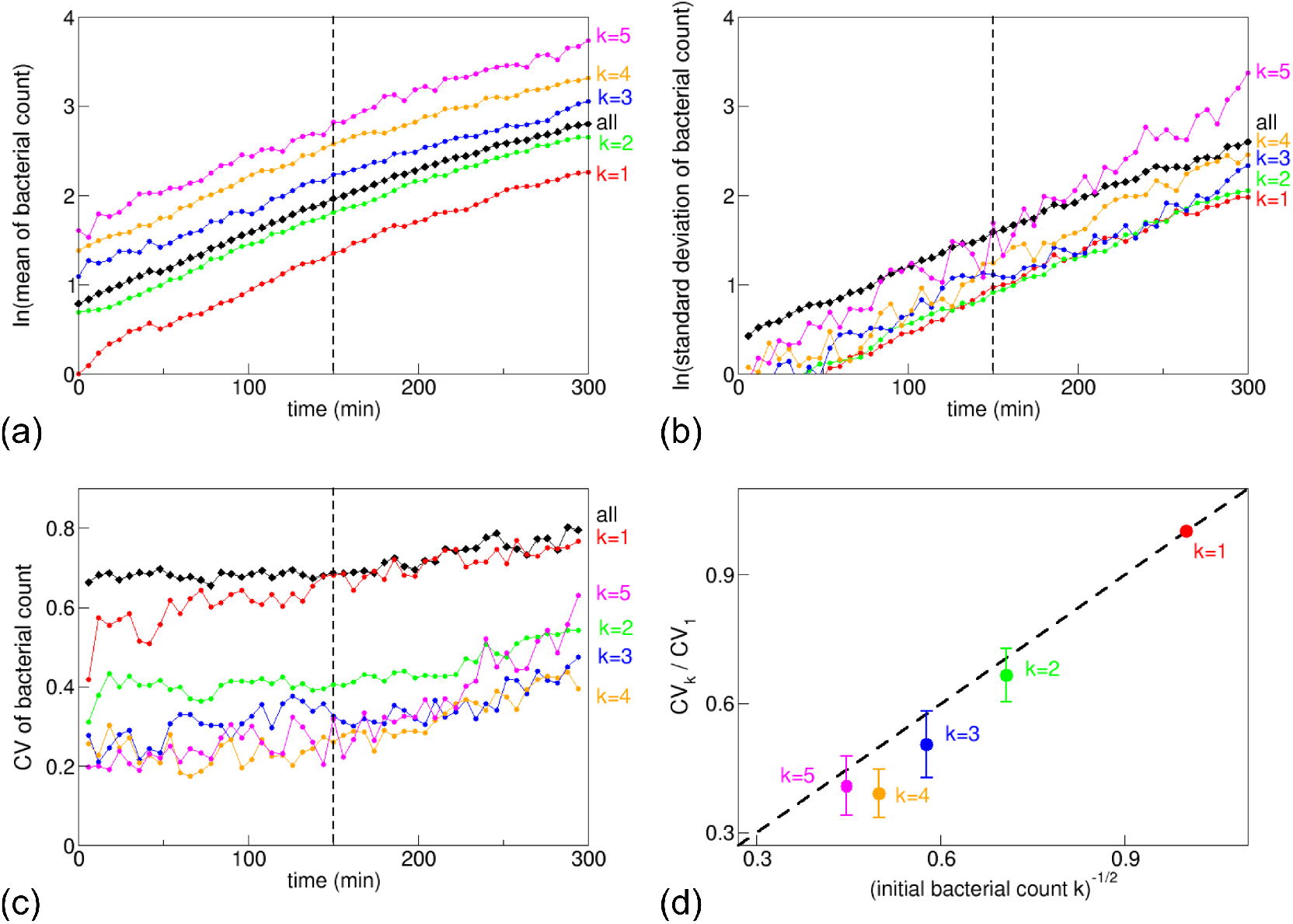
Stochastic growth of encapsulated *E. coli*; comparison to the predictions of the Bellman-Harris model [46]. (a): The mean bacterial count grows exponentially in time, for the entire collection of non-empty droplets (black data), and for the sub-groups of droplets, sorted by initial bacterial count *k* (red: *k* = 1, green: *k* = 2, blue: *k* = 3, orange: *k* = 4, magenta: *k* = 5. The vertical dashed line indicates 150min; the growth rate was obtained by fitting to the data for 0-150min. (b): The standard deviation of the bacterial count also grows exponentially in time (colours as in (a). (c): The coefficient of variation (CV = standard deviation / mean) is approximately constant at early times (colours as in (a)). The values for CV reported in the text were computed by averaging over the data for 0-150min. (d): The coefficient of variation *CV*_*k*_ for droplets starting with *k* bacteria scales with 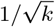, as predicted by Barizien *et al*.’s generalisation of the Bellman-Harris theory [22].

#### Inferring single-cell division time statistics

Using the BH model, we can also infer the mean *τ* and standard deviation *σ* [22]) of single-cell division times. Assuming that the single-cell division times are Gaussian distributed, Barizien et al. showed that [22]

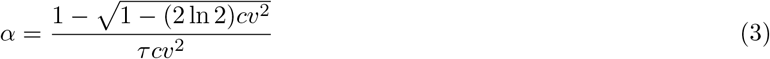

and

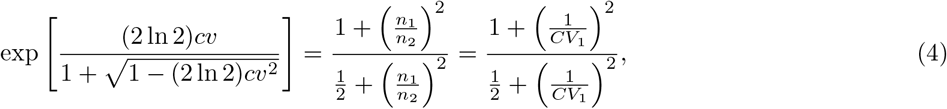

where *cv* = *σ/τ* is the coefficient of variation of the single-cell division-time distribution. Using our 68 trajectories that started with a single bacterium, we obtain *α* = 0.00777 ± 0.00007min^−1^, from a linear fit of the log-mean population growth over the first 150min (Figure 5(a), red data points). Using also our result that *CV*_1_ = 0.60 ± 0.05 (averaged over the first 150min), we can solve numerically Eqs (3) and (4) to estimate the single-cell division time parameters: *τ* = 86 ± 2min and *cv* = 0.20 ± 0.03. Single-cell division statistics can also be measured directly, for example, in the mother machine [51]. While the reported values for *cv* vary [43, 52, 53], and depend on conditions such as growth medium [53], our result is close to that of *cv* = 0.24 reported by Wallden et al. [53] for mother machine measurements, for bacteria that were growing at a similar rate to ours (mean growth rate 0.011min^−1^).

## Discussion

In this paper, we have presented a microfluidic system, with image analysis protocol, that allows direct counting of individual bacteria during growth in microfluidic-produced emulsion droplets. While a number of microfluidic devices have been developed previously to monitor bacterial growth in droplets [6, 11, 17, 18, 20–22, 33], these have mostly used the total fluorescence of a droplet as a proxy for population size. The few cases where bacterial numbers have been directly counted have not allowed high-throughput counting [19]. To our knowledge, our study is the first to achieve automated direct counting of individual bacteria in a high-throughput manner. Direct counting avoids technical issues caused by fluorescence heterogeneity between bacteria [34] and changes in bacterial fluorescence intensity during growth [35]. Direct counting also removes the need for the use of advanced fluorescence background subtraction methods [22]. Furthermore, knowledge of the absolute population size is important when comparing experimental results to theory for stochastic population growth [22].

Using our system, we analysed in detail the population size statistics for 188 non-empty droplets containing *E. coli* growing on minimal media. Extending the earlier analysis of Barizien *et al*. (who did not have access to absolute bacterial counts), we found excellent agreement between our data and the predictions of the Bellman-Harris model [46], which assumes that cell division events are independent, with the division times being drawn from a fixed distribution. Under the assumption that the distribution of single-cell division times is Gaussian [22], we could infer from our population growth trajectories the mean and coefficient of variation of the single-cell division times. The fact that we obtain good fits to the memory-less Bellman-Harris model is consistent with mother machine observations that *E. coli* shows little inheritance of growth rate under standard, aerobic growth conditions [51]. However there is evidence that growth rate can be more strongly inherited under other conditions, such as anaerobic growth [54]. It would therefore be interesting to investigate population growth statistics for bacteria in droplets under a wide range of conditions, as well as to investigate the power of such data to discriminate between alternative population growth models.

Our system comprises two parts, the first of which generates a microfluidic droplet emulsion whilst the second acts as a reservoir for imaging and analysing droplet-based bacterial populations. Crucially, the reservoir height is small (11*µ*m), to prevent bacteria from leaving the focal plane during imaging. Because droplet generation is more robust and reliable in a deeper device, we have fabricated the two parts of the device with different heights, and connected them via a short piece of tubing. This does, however, introduce some delay between droplet creation and imaging (around 90 mins in total to transfer droplets and fill the reservoir). In future we plan to eliminate this delay by integrating the two components into a single multilevel device.

Image processing is an important aspect of our study. The image processing algorithm must be sophisticated enough to track hundreds of droplets and count individual bacterial cells within them. The algorithm must also be efficient enough to process large volumes of data within a reasonable time frame. The current version of our image processing algorithm runs on a standard desktop computer with a typical experimental run taking less than 4 days to fully process on a standard desktop PC (including manual counting for optimisation of the binarisation threshold, manual stitching of fields of view and and manual image preprocessing). We are currently investigating options for parallelising the code to improve image processing speed, and automating the steps that are currently done by hand, such as stitching together of multiple fields of view. Other parts of the image processing algorithm that could be made more sophisticated, or automated, include the droplet identification and determining the optimal threshold for binarising the images prior to counting of the bacteria.

In this work, we chose to use a 6 minute time interval between successive images of the same field of view. This was the shortest time interval that allowed all 18×14 fields of view in the grid to be imaged in both brightfield and fluorescence, before a new round of imaging began. This time interval is substantially smaller than the expected doubling time of the *E. coli* bacteria used, and allowed good tracking of droplets between frames. However, the 6 minute time interval is too long to allow dynamical tracking of individual bacteria within droplets, which might be interesting for some applications, such as studies of bacterial motility [55]. Recording of short movies of a given field of a view, rather than static images, might be a fruitful direction should dynamical information be required.

Our approach to bacterial counting relies, at present, on fluorescence images – therefore we have used a bacterial strain that expresses a fluorescent protein. Fluorescent strains of bacteria are widespread, and for applications with non-fluorescent bacteria one could perhaps use a fluorescent dye. However in future we hope also to develop bacterial counting via segmentation of phase contrast images, using more sophisticated image processing techniques.

In our experiments, we observe that our strain of *E. coli* tends to form clusters in the droplets at long times, which prevents the counting algorithm from working. This could perhaps be avoided by using a non-aggregating bacterial strain (e.g. a strain that does not produce appendages such as fimbriae that are implicated in aggregation). However, bacterial clustering is of biological interest in itself, for example as a first stage in biofilm formation [56, 57]. Therefore it might be interesting to develop image analysis algorithms that could detect and analyse the onset of clustering in microfluidic droplets, perhaps via the observed fluctuations in the bacterial count.

It is important to note that the approach presented here allows for retrospective investigation of phenomena occurring in a given droplet. For example, if a growth trajectory shows unexpected dynamics, one can extract the image series corresponding to that droplet and inspect the images to see what has happened. This provides a useful way to control for errors (e.g. in bacterial counting) but also to investigate biological phenomena such as the onset of bacterial aggregation.

Future improvements to our device may also include the use of injection to allow chemical perturbations to droplets after they are created (e.g. the injection of other bacterial species to test for production of potential biopharmaceuticals [11]), and the use of droplet sorting [58] to allow the extraction and further processing of droplets with interesting properties, for example for biotechnological applications.

## Conclusions

We developed a microfluidic device and image analysis protocol that can count individual bacteria during growth in microfluidic droplets. This allows high throughout tracking of the dynamics of small bacterial populations, without the need for proxy measures of population size such as integrated droplet fluorescence. Analysing growth trajectories for multiple small populations of *E. coli* bacteria we find good agreement with the Bellman-Harris model, which assumes that individual-cell lifetimes are independent. We are also able to infer parameters of the single-cell lifetime distribution from population trajectories. We hope that our approach will contribute to the numerous fundamental and biotechnological applications of droplet-based bacterial culture.

## Materials and Methods

### 1. Microfluidic device fabrication

Our PDMS microfluidic device was created as follows. Microfluidic master moulds with depths of 34.5*µ*m and 11.5*µ*m (for the droplet generator and reservoir respectively) were fabricated at the Scottish Microelectronics Centre using standard methods. SU8-3010 was spin coated on a 4′′ silicon wafer to the required thickness (2min at 2000 rpm for the 34.5*µ*m-depth and 2min at 4000rpm for the 11.5*µ*m depth). The wafers were baked (1min at 65°C followed by 10min at 95°C), exposed to UV light through a photomask (2min) then baked again (1min at 65°C then 2min at 95°C) and finally rinsed for 4min in propylene glycol methyl ether acetate (PGMEA). We used a 4′′ low reflective chrome photomask printed onto a 1.5mm soda lime glass base substrate at a resolution of 128k DPI (JD-PhotoData).

To make a device, 30g of PDMS was prepared by mixing 10 parts elastomer to 1 part crosslinking agent for 5min. Air bubbles were removed by exposing to vacuum (30min). The PDMS was then poured over the microfluidic master to a depth of 15mm then placed under vacuum to remove air bubbles, before baking (90min at 80°C). Excess PDMS was removed and inlet and outlet holes were created using a biopsy punch. 20mm-length glass capillaries were inserted into the inlet and outlet holes and flushed with compressed air to remove debris. The PDMS surface was cleaned using compressed air and repeated gentle application of adhesive tape.

The PDMS device was plasma treated, together with a glass microscope slide (Diener Zepto plasma oven, oxygen plasma for 55s at 35W), and the PDMS and glass were bonded together by gentle pressing. Both the droplet generator and the reservoir were mounted on the same microscope slide. The glass capillaries were then sealed to the PDMS with a 2-part epoxy resin. The tube connecting the generator and reservoir was connected after the expoxy had dried. Finally 500*µ*l of PicoGlide (Dolomite Microfluidics) was injected into the device with a 1ml syringe and left for 60min before flushing. PicoGlide contains a fluorinated polyether polymer that forms a covalently-bonded layer on the PDMS and glass surfaces, enhancing droplet stability [www.spherefluidics.com].

### 2. Bacterial strains and media

Our experiments used *E. coli* strain RJA002 [50]. RJA002 is a derivative of strain MG1655, with a chromosomal insertion of a gene encoding yellow fluorescence protein (YFP) under the control of the phage lambda P1 promoter, which is constitutively expressed. RJA002 also contains a chloramphenicol resistance cassette. It was created by P1 transduction from strain MRR, supplied by the Elowitz lab [34, 50].

All experiments were performed in M9 minimal medium, supplemented with 0.4%w/v glucose. The medium was prepared in-house; briefly, 100ml of M9 X4 salt stock (28g Na_2_HPO_4_, 12g KH_2_PO_4_, 2g NaCl and 4g NH_4_Cl in 1l sterile H_2_O) was mixed with 300ml sterile H_2_O, 800*µ*l of 1M MgSO_4_ and 40*µ*l of 1M CaCl_2_, then 8ml of 20%w/v glucose was added.

### 3. Device setup

Our experimental protocol was as follows. An overnight culture of *E. coli* strain RJA002 was prepared in M9 minimal medium. This was used to inoculate a fresh culture 1:400 which was incubated (37°C, 200rpm) for 1.5h before encapsulation into droplets.

For droplet creation, the device was first filled with the oil phase (FC40 containing 2.5%w/w PicoSurf) using a syringe pump (New Era NE-1002X) at 50*µ*l h^−1^. A second, identical, syringe pump was then used to flow the aqueous phase (bacterial suspension) into the device at 50*µ*l h^−1^, and the device was immersed in water at 37°C using the bespoke microscope-mounted chamber.

Once both the aqueous and oil phase had reached the flow-focusing junction and droplet generation had begun, the device was allowed to equilibrate for 15min, then the flow rates of the aqueous and oil phases were adjusted to 20*µ*l h^−1^ and 80*µ*l h^−1^ respectively. After a further 15min the outlet tubing was disconnected so that the droplet reservoir began to fill – a process that took 1.5-2h. Once the reservoir was full with monodisperse droplets, the syringe pumps were stopped and the inlet and bridge tubing were severed, arresting droplet movement within the reservoir. Image acquisition then commenced. During the entire 8h period of image acquisition the temperature of the water in which the device was immersed was maintained at 37°C.

### 4. Microscopy

Our experiments were performed using a Nikon Eclipse Ti epi-fluorescence inverted microscope with a Nikon Plan Fluor 20x/0.5 objective. We used a Hamamatsu Orca-Flash4.0 V2 (C11440-22CU) camera with a CMOS image sensor with 6.5*µ*m x 6.5*µ*m pixel size and 2048 × 2048 imaging pixels. Fluorescence imaging used a broad-spectrum Mercury lamp with a Chroma EYFP filter. The brighteld channel exposure was set to 10ms, the fluorescence exposure was set to 50ms and we used 2×2 pixel binning in both channels. Automatic focussing (perfect focus) was used, with the focus being initially adjusted manually for optimal imaging of bacteria viewed in the uorescence channel. The reservoir was imaged by scanning a grid, typically consisting of 18×14 fields of view in the brightfield and fluorescence channels. The whole microfluidic reservoir was imaged once per 6min and a full experimental run was 8h long. Therefore, each grid position was imaged 80 times.

### 5. Image analysis: droplet detection and tracking

Droplets were detected using Matlab’s `imfindcircles` function, which searches for circular objects in an image using a circular Hough transform. The function outputs a data array of centre positions and radii for all circular objects found in the image. The sensitivity factor and edge threshold value were adjusted manually to optimise droplet detection. The detected circles were filtered by a minimum and maximum radius, allowing for the exclusion of rare non-monodisperse droplets and circular features in the microfluidic device structure, such as reservoir supports.

Droplet tracking was performed using an adapted particle tracking algorithm [49]. This algorithm is based on the Matlab `track` function, which considers all possible identifications of the old positions with the new positions, and chooses the identification which results in the minimal total squared displacement. Droplet tracking works well in our system because very few droplets move by more than their own radius during the 6min interval between images. The resulting data array contains an identification number for each droplet that allows it to be followed in time. We note that in this paper we only analyse trajectories for droplets that were tracked over the entire duration of the experiment.

### 6. Image analysis: binarization threshold optimisation

Binarization of the fluorescence images is a key step in our bacterial counting algorithm. This amounts to setting every pixel to either ‘on’ or ‘off’, depending on whether the pixel intensity is above or below a threshold value. To optimise the binarization threshold, we select images corresponding to 4 fields of view, distributed across the reservoir, for a timepoint midway through the experiment. We perform manual counts of the number of bacteria within each droplet, using the fluorescence images overlaid by the droplet boundaries. We then binarize the images using a given threshold and quantify the number of distinct objects (defined as pixel islands that are not 8-connected) within each droplet boundary. This procedure is performed for a range of binarization thresholds and the sum of the absolute differences between the manual and automated counts is plotted as a function of the binarization threshold. The optimal threshold value is taken as the minimum of this curve (Figure 2(c)).

### 7. Handling large image datasets

Each experiment results in a large volume of image data, such that the stitched 18×14 grid of images is too large to be processed on a standard desktop PC. Therefore in practice we split the grid into a small number of ‘subsections’ (typically 9), that are processed separately, but using the same image processing parameters (e.g. optimal binarization threshold). This allows the data to be processed on a standard desktop PC. Splitting the data in this way could introduce some errors due to some droplets being cut in half at the edge of a subsection - although the boundaries between subsections were chosen to avoid this as much as possible. These errors should be caught by the manual image checking step in the data processing.

### 8. Data analysis for Poisson encapsulation of encapsulated E. coli

To analyse the distribution of bacterial counts in the first set of images taken in our experiments, we first fitted to a Poisson distribution *p*_*λ*_(*N*) = *λ*^*k*^*e*^−*λ*^*/k*!, where *N* is the bacterial count measured in a given droplet and *λ* is the Poisson parameter (to be fitted). This resulted in poor agreement with the data (Figure 4(a); red shaded area). We then performed computer simulations in which initially Poisson-distributed populations of bacteria were assumed to grow exponentially at rate *µ* for time *τ*. This growth was assumed to be deterministic (i.e. we neglect the inherent stochasticity of bacterial growth). The simulated distribution of final bacterial counts was then fitted to our experimental data, with the fit parameters being the Poisson parameter *λ*, and the product *µτ* (since only the product is relevant for exponential growth). This produced a much better fit (Figure 4(a), green shaded area). The best fit parameters for our experimental dataset were *λ* = 0.77, and *µτ* = 0.36. The fitted value for *µτ* is about half the expected value given the known wait time and growth rate (95min × 0.0077 min^−1^ = 0.73). This might be accounted for if some bacteria were still in the lag phase of growth when the droplets were created; in future experiments we plan to ensure a longer period of exponential growth prior to droplet encapsulation.

### 9. Dataset for analysis of stochastic growth of E. coli

For the analysis presented in Figure 5, we used a dataset consisting of 375 droplets, containing *E. coli* strain RJA002. Of these droplets, 184 were empty, *i*.*e*. they had a zero bacterial count at the start. To avoid artefacts that can skew the bacterial number distribution, we manually checked the microscopy images for all droplets that started with non-zero counts but had counts less than 7 bacteria the end of the 8h experimental run. This resulted in the exclusion of 3 droplets in which small pieces of dust had been incorrectly counted as bacteria. Therefore our final dataset consisted of 188 non-empty droplet trajectories. In our analysis, the coefficients of variation *CV*_*k*_ were computed by averaging over the first 150min of our dataset (omitting the first datapoint for which by definition the standard deviation of the trajectories is zero). Error bars in the reported values of *CV*_*k*_ refer to 95% confidence intervals, obtained by bootstrapping the datasets.

## Acknowledgments

We thank Patrick Warren (Unilever R&D, Port Sunlight) for support, guidance, and comments on the manuscript. We also acknowledge valuable discussions with Aidan Brown, Stefano Pagliara and Bartek Waclaw. We thank Angela Dawson for her assistance with microbiological techniques and strains, and Louis Berridge for assistance with MatLab. DT and NV were supported by the EPSRC CDT on “Soft and Functional Interfaces” (SOFI). RJA was supported by a Royal Society University Research Fellowship and by the European Research Council under Consolidator grant 682237 EVOSTRUC.

## References

[1] A. B. Theberge, F. Courtois, Y. Schaerli, M. Fischlechner, C. Abell, F. Hollfelder, and W. T. Huck, Ang. Chemie Int. Ed. 49, 5846 (2010).

[2] J. J. Agresti, E. Antipov, A. R. Abate, K. Ahn, A. C. Rowat, J.-C. Baret, M. Marquez, A. M. Klibanov, A. D. Griffiths, and D. A. Weitz, Proc. Natl. Acad. Sci. USA 107, 4004 (2010).

[3] O. J. Miller, A. El Harrak, T. Mageat, J.-C. Baret, L. Frenz, B. El Debs, E. Mayot, M. L. Samuels, E. K. Rooney, P. Dieu, et al., Proc. Natl. Acad. Sci. USA 109, 378 (2012).

[4] S. Köster, F. E. Angilè, H. Duan, J. J. Agresti, A. Wintner, C. Schmitz, A. C. Rowat, C. A. Merten, D. Pisignano, A. D. Griffiths, et al., Lab Chip 8, 1110 (2008).

[5] T. S. Kaminski, O. Scheler, and P. Garstecki, Lab Chip 16, 2168 (2016).

[6] K. Leung, H. Zahn, T. Leaver, K. M. Konwar, N. W. Hanson, A. P. Pagé, C.-C. Lo, P. S. Chain, S. J. Hallam, and C. L. Hansen, Proc. Natl. Acad. Sci. USA 109, 7665 (2012).

[7] J.-U. Shim, L. F. Olguin, G. Whyte, D. Scott, A. Babtie, C. Abell, W. T. S. Huck, and F. Hollfelder, J. Am. Chem. Soc. 131, 15251 (2009).

[8] Y. Zhang, Y. P. Ho, Y. L. Chiu, H. F. Chan, B. Chlebina, T. Schuhmann, L. You, and K. W. Leong, Biomaterials 34, 4564 (2013).

[9] S. Abalde-Cela, A. Gould, X. Liu, E. Kazamia, A. G. Smith, and C. Abell, J. R. Soc. Interface 12, 20150216 (2015).

[10] N. Shembekar, H. Hu, D. Eustace, and C. A. Merten, Cell Rep. 22, 2206 (2018).

[11] L. Mahler, K. Wink, R. Beulig, K. Scherlach, M. Tovar, E. Zang, K. Martin, C. Hertweck, D. Belder, and M. Roth, Sci. Rep. 8, 13087 (2018).

[12] J. Q. Boedicker, M. E. Vincent, and R. F. Ismagilov, Angew. Chemie Int. Ed. 48, 5908 (2009).

[13] A. B. Chang, J. N. Wilking, S.-H. Kim, H. C. Shum, and D. A. Weitz, Small 11, 3954 (2015).

[14] J. Park, A. Kerner, M. A. Burns, and X. N. Lin, PLoS ONE 6, e17019 (2011).

[15] X. Guo, K. P. T. Silva, and J. Q. Boedicker, Phys. Biol. 16, 036001 (2019).

[16] Y. Bai, S. N. Patil, S. D. Bowden, S. D. Poulter, J. Pan, G. P. Salmond, M. Welch, W. S. Huck, and C. Abell, Int. J. Mol. Sci. 14, 10570 (2013).

[17] W. Postek, P. Gargulinski, O. Scheler, T. S. Kaminski, and P. Garstecki, Lab Chip 18, 3668 (2018).

[18] F. Lyu, M. Pan, S. Patil, J.-H. Wang, A. Matin, J. R. Andrews, and S. K. Tang, Sensors Actuator B 270, 396 (2018).

[19] P. Sabhachandani, S. Sarkar, P. C. Zucchi, B. A. Whitfield, J. E. Kirby, E. B. Hirsch, and T. Konry, Microchimica Acta 184, 4619 (2017).

[20] A. M. Kaushik, K. Hsieh, L. Chen, D. J. Hin, and J. C. Liao, Biosensors Bioelectronics 97, 260 (2017).

[21] O. Scheler, K. Makuch, P. R. Debski, M. Horka, A. Ruszczak, N. Pacocha, K. Sozański, O. Smolander, W. Postek, and P. Garstecki, Sci. Rep. 10, 3282 (2020).

[22] A. Barizien, M. S. Suryateja Jammalamadaka, G. Amselem, and C. N. Baroud, J. R. Soc. Interface 16, 20180935 (2019).

[23] J. Pan, A. L. Stephenson, E. Kazamia, W. S. Huck, J. S. Dennis, A. G. Smith, and C. Abell, Integr. Biol. 3, 1043 (2011).

[24] S. P. Damodaran, S. Eberhard, L. Boitard, J. G. Rodriguez, Y. Wang, N. Bremond, J. Baudry, J. Bibette, and F. A. Wollnam, PLoS ONE 10, e01118987 (2015).

[25] D. Cottinet, F. Condamine, N. Bremond, A. D. Griffiths, P. B. Rainey, J. A. G. M. de Visser, J. Baudry, and J. Bibette, PLoS ONE 11, e0152395 (2016).

[26] D. J. Collins, A. Neild, A. DeMello, A.-Q. Liu, and Y. Ai, Lab Chip 15, 3439 (2015).

[27] E. W. M. Kemna, R. M. Schoeman, F. Wolbers, I. Vermes, D. A. Weitz, and A. van den Berg, Lab Chip 12, 2881 (2012).

[28] Y.-J. Eun, A. S. Utada, M. F. Copeland, S. Takeuchi, and D. B. Weibel, ACS Chem. Biol. 6, 260 (2011).

[29] S. Jakiela, T. S. Kaminski, O. Cybulski, D. B. Weibel, and P. Garstecki, Angew. Chem. Int. Ed. 52, 8908 (2013).

[30] A. Grodrian, J. Metze, T. Henkel, K. Martin, M. Roth, and J. M. Köhler, Biosensors and Bioelectronics 19, 1421 (2004).

[31] R. Ramji, M. Wang, A. A. S. Bhagat, D. T. S. Weng, N. V. Thakor, C. T. Lim, and C.-H. Chen, Biomicrofluidics 8, 034104 (2014).

[32] R. J. Best, J. J. Lyczakowski, S. Abalde-Cela, Z. Yu, C. Abell, and A. G. Smith, Anal. Chem. 88, 10445 (2016).

[33] S. Huang, J. K. Srimani, A. J. Lee, A. J. Lopatkin, K. W. Leong, and L. You, Biomaterials 61, 239 (2015).

[34] M. B. Elowitz, A. J. Levine, E. D. Siggia, and P. S. Swain, Science 297, 1183 (2002).

[35] O. Gefen, O. Fridman, I. Ronin, and N. Q. Balaban, Proc. Natl. Acad. Sci. USA 111, 556 (2014).

[36] K. Stevenson, A. McVey, I. Clark, P. Swain, and T. Pilizota, Sci. Rep. 6, 38828 (2016).

[37] R. J. Allen and B. Waclaw, Rep. Prog. Phys. 82, 016601 (2019).

[38] H. Lu, O. Caen, J. Vrignon, E. Zonta, Z. El Harrak, P. Nizard, J. Baret, and V. Taly, Sci. Rep. 7, 1366 (2017).

[39] N. Q. Balaban, J. Merrin, R. Chait, L. Kowalik, and S. Leibler, Science 305, 1622 (2004).

[40] J. B. Deris, M. Kim, Z. Zhang, H. Okano, R. Hermsen, A. Groisman, and T. Hwa, Science 342, 1237435 (2013).

[41] J. Coates, B. R. Park, D. Le, E. Şimşek, W. Chaudry, and M. Kim, eLife 7, e32976 (2018).

[42] Y. Wakamoto, N. Dhar, R. Chait, K. Schneider, F. Signorino-Gelo, L. S., and J. D. McKinney, Science 339, 91 (2013).

[43] S. Taheri-Araghi, S. Bradde, M. Vergassola, S. Jun, J. T. Sauls, N. Hill, P. A. Levin, and J. Paulsson, Curr. Biol. 25, 385 (2015).

[44] J. Lin and A. Amir, Cell Syst. 5, 1 (2017).

[45] M. Hashimoto, T. Nozoe, H. Nakaoka, R. Okura, S. Akiyoshi, K. Kaneko, and Y. Wakamoto, Proc. Natl Acad. Sci. USA 113, 3251 (2016).

[46] R. Bellman and T. Harris, Ann. Math. 55, 385 (1952).

[47] P. Garstecki, I. Gitlin, W. DiLuzio, G. M. Whitesides, E. Kumacheva, and H. A. Stone, Applied Physics Letters 85, 2649 (2004).

[48] J. Shemesh, T. Ben Arye, J. Avesar, J. H. Kang, A. Fine, M. Super, A. Meller, D. E. Ingber, and S. Levenberg, Proc. Natl. Acad. Sci. USA 111, 11293 (2014).

[49] J. C. Crocker and D. G. Grier, J. Colloid Interface Science 179, 298 (1996).

[50] D. P. Lloyd, Ph.D. thesis (2015), URL http://hdl.handle.net/1842/10509.

[51] P. Wang, L. Robert, J. Pelletier, W. L. Dang, F. Taddei, A. Wright, and S. Jun, Current Biology 20, 1099 (2010), ISSN 0960-9822, URL https://www.sciencedirect.com/science/article/pii/S0960982210005245?via{\%}3Dihub.

[52] M. Campos, I. V. Surovtsev, S. Kato, A. Paintdakhi, B. Beltran, S. E. Ebmeier, and C. Jacobs-Wagner, Cell 159, 1433 (2014).

[53] M. Wallden, D. Fange, E. G. Lundius, Ö. Baltekin, and J. Elf, Cell 166, 729 (2016).

[54] W. H. Hughes, J. gen. Microbiol. 12, year = 1955, 265 (????).

[55] I. D. Vladescu, E. J. Marsden, J. Schwarz-Linek, V. A. Martinez, J. Arlt, A. N. Morozov, D. Marenduzzo, M. E. Cates, and W. C. K. Poon, Phys. Rev. Lett. 113 (2014).

[56] K. Kragh, J. Hutchison, G. Melaugh, C. Rodesney, A. Roberts, Y. Irie, P. Jensen, S. Diggle, R. Allen, V. Gordon, et al., mBio 7, e00237 (2018).

[57] G. Melaugh, J. Hutchison, K. Kragh, Y. Irie, A. Roberts, T. Bjarnsholt, S. Diggle, V. Gordon, and R. Allen, PLoS ONE 11, e0149683 (2016).

[58] J.-C. Baret, O. J. Miller, V. Taly, M. Ryckelynck, A. El-Harrak, L. Frenz, C. Rick, M. L. Samuels, J. B. Hutchison, J. J. Agresti, et al., Lab on a Chip 9, 1850 (2009).

[59] We dilute the as-supplied 5%w/w in FC40 solution (https://spherefluidics.com/store/-surf-1-5-w-w-in-fc-40/?v=79cba1185463) by a factor of 2 with FC40; Pico-Surf is widely used in microfluidic droplet applications - see the website for example application notes.

