## supplementary material for "Tracking the stochastic growth of bacterial populations in microfluidic droplets"

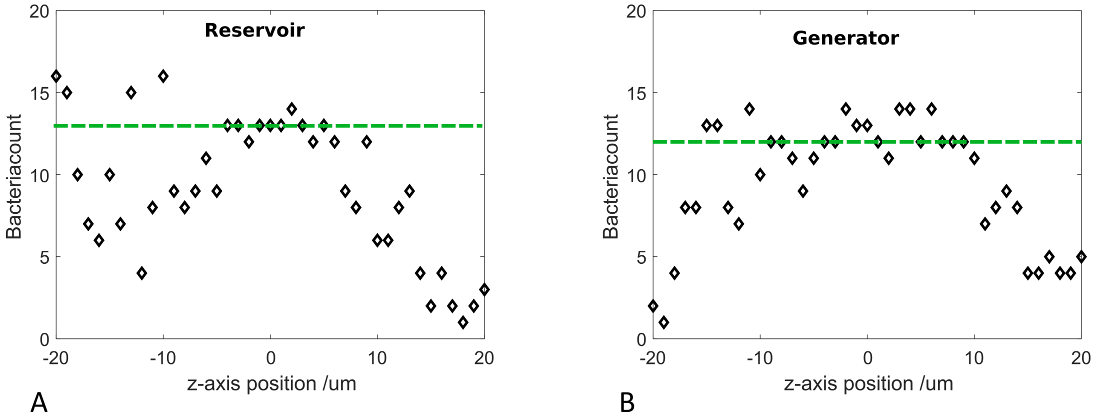


FIG. S1: **Squashing droplets in the reservoir keeps bacteria in focus**. In our device, the reservoir height is limited to 11.5$\mu$m, in order to squash droplets vertically and prevent bacteria from leaving the focal plane. To test whether bacteria indeed remain in the focal plane in our reservoir, we analysed in detail how the quality of detection of single cells in our device changes as we scan vertically through the device. The plots show the bacterial count, as the z position is scanned, for a single field of view in the reservoir part of the device (A), compared with a single field of view in the generator part of the device (B). The counts are summed over all droplets in the field of view (12 droplets for the reservoir A, and 5 droplets for the generator B). Manual counts (shown by the dashed line) indicated that the correct bacterial counts were 13 (reservoir, A) and 12 (generator, B) - correct counts are indicated by the green dashed line. In the reservoir, where the droplets are squashed, there is a clear z-range where the bacterial count is constant and correct; the width of this range is approximately 12$\mu$m, implying that we can consistently image all bacteria within the squashed droplets in the reservoir of height 11.5$\mu$m. Outside this range, the count decreases sharply since the droplet is no longer in the focal plane. In contrast, for the field of view in the generator part of the device (B), which has height 35.5$\mu$m, implying that the droplets are less squashed, the bacterial count varies more, even over the region where the droplets are in focus. This is probably because bacteria are able to move in and out of the focal plane in these less squashed droplets. For both the reservoir and generator images, the image analysis parameters were optimised for the image corresponding to the central (focussed) z-position (z=0).

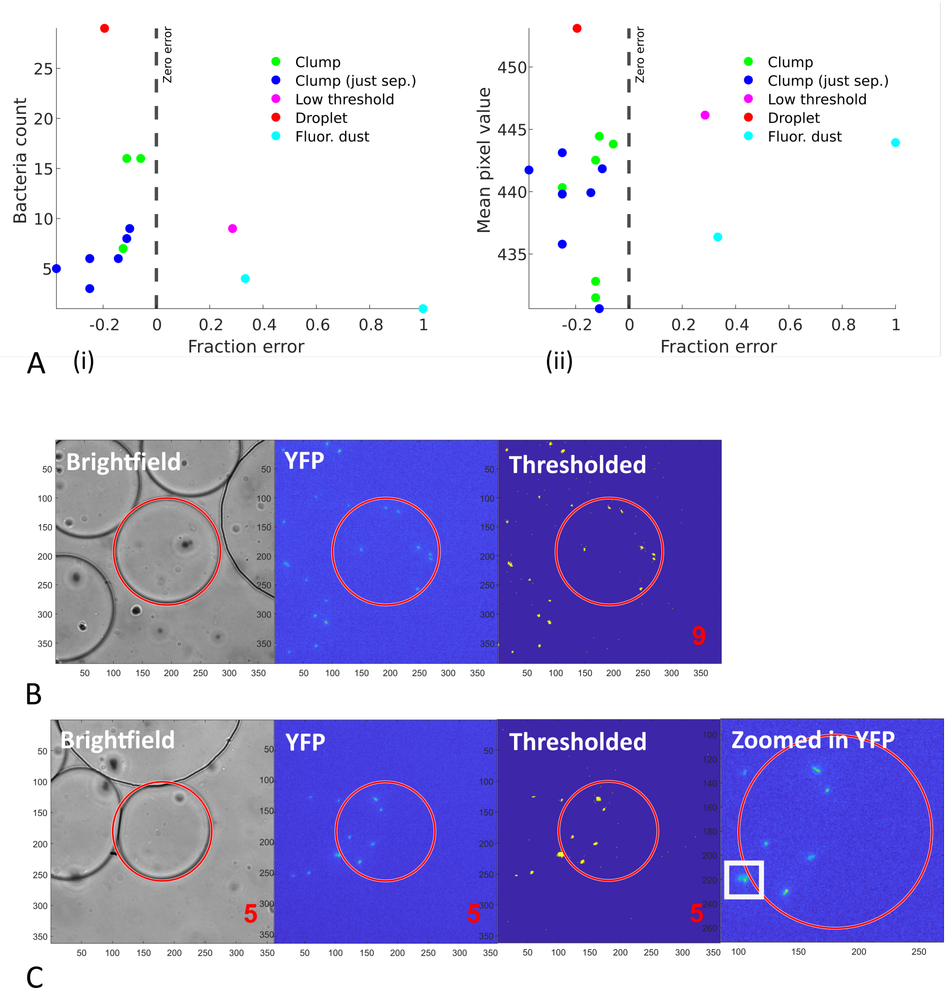


FIG. S2: **Analysis of counting errors**. To understand the origin of errors in droplet counting, we analysed a dataset consisting of 118 droplets, with a total of 475 bacteria, imaged after 120 mins of growth. These droplets were manually counted and the results compared with the output of our counting algorithm. We found 17 droplets for which the automated bacterial count differed from the manual count. For these droplets, panel A investigates whether the count error correlates with either (i) the number of bacteria in the droplets (counted manually), or (ii) the total droplet fluorescence. We did not find any such correlation, suggesting that there is not a simple link between bacterial density and count error, for early-time trajectories. Next, we looked in detail at the images for these 17 droplets, and categorized the errors into 5 types. 6/17 droplets showed clumping of bacteria, causing multiple bacteria to be counted as one unit. 7/17 droplets showed dividing cells, where the two daughter cells have just separated but are counted by the algorithm as one unit. 1/17 droplets had a too low threshold, so that random noise pixels were counted as bacteria. 1/17 droplets had an incorrect droplet boundary position, so that bacteria near the edge of the droplet are not counted. 2/17 droplets contained fluorescent dust particles that were incorrectly counted as bacteria. The data points in panel A are colour-coded according to the source of the error (see legend). Panel B shows images (brightfield, YFP fluorescence and the binary thresholded fluorescence image) for the droplet that had a too low threshold value, so that random pixels are counted as bacteria. Here, the correct droplet count is 7 but the algorithm counts 9 bacteria. Panel C shows an image with 3 issues: an incorrect droplet boundary, some fluorescent dust (shown in the box in the zoomed in image), and a recently divided pair of daughter bacteria that are counted as one.

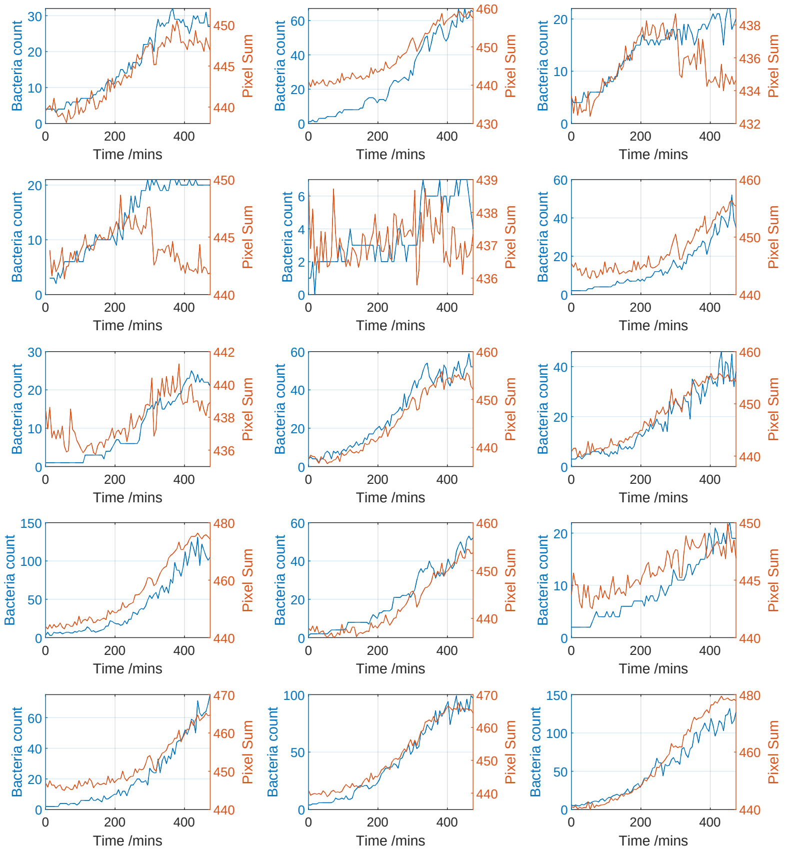


FIG. S3: **Comparing total fluorescence with automated bacterial counts**. Trajectories of total fluorescence (pixel sum normalised by droplet area) and of bacterial counts (obtained using our automated algorithm), are shown for 15 example droplets, over the course of a 480 min growth experiment in M9 glucose minimal medium (see Methods). The bacterial counts are shown in blue (left hand axis) while the integrated fluorescence is shown in orange (right hand axis). These droplets correspond to a subset of those analysed in Figure 3(b) of the main text. In most droplets, the total fluorescence and bacterial counts follow the same trend; however in some droplets total fluorescence decreases at late times while bacterial count levels off but does not decrease. This may be due to decreased fluorescence of individual bacteria as they enter stationary phase.

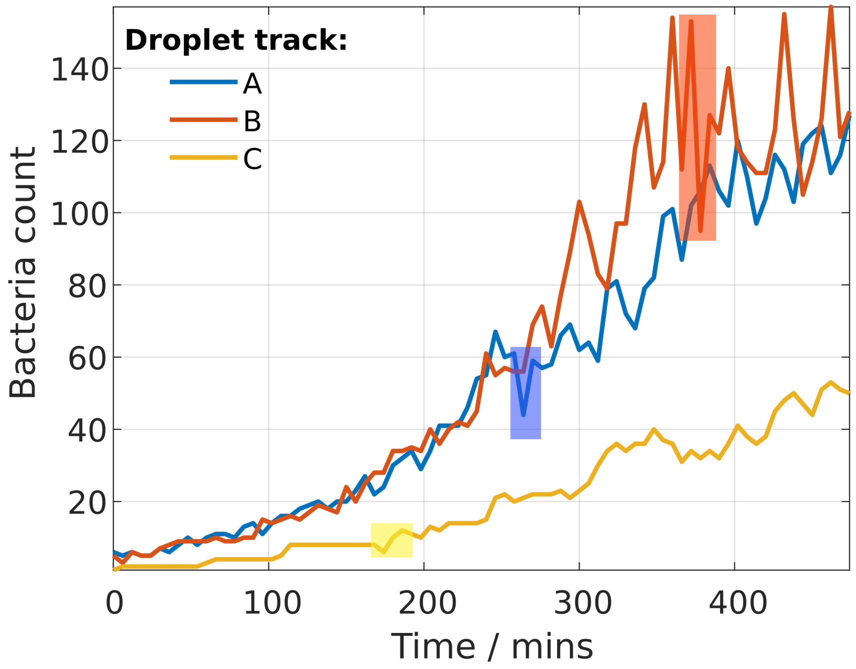


FIG. S4: **Transient dips in the bacterial count during growth trajectories**. We selected three droplet trajectories (bacterial count vs time) which showed transient dips in the bacterial count. Since we do not expect to see bacterial death under these experimental conditions, we hypothesized that these dips were caused by counting errors. In the three selected trajectories, the dips in the bacterial count are indicated by the shaded boxes. The microscopy images corresponding to these sections of the trajectories are investigated in the following figures.

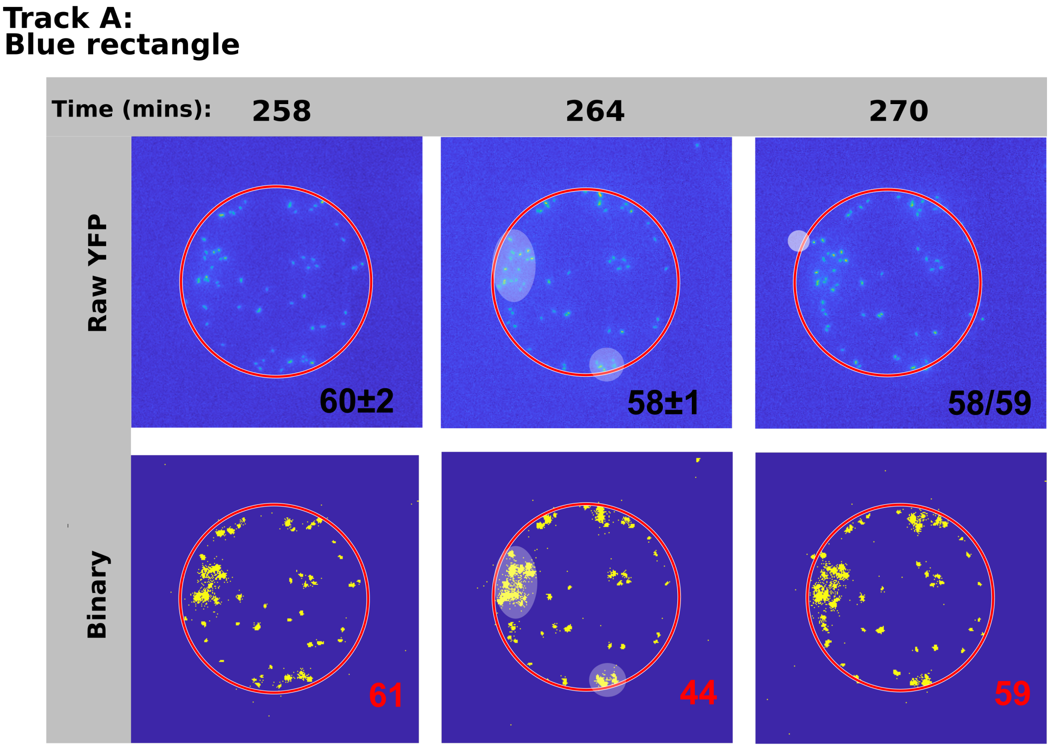


FIG. S5: **Transient dip: example A**. Microscopy images (fluorescence image and binary thresholded image) from the trajectory shown in blue in Figure S4. The images correspond to the times leading up to the dip in bacterial count at 270min (shaded blue area in Figure S4). The numbers on the images show the bacterial counts (black numbers are manual counts; red numbers are automated counts). The red circle shows the droplet boundary as detected by our software. At 264min (middle images), several bacteria become close together and are counted as one, resulting in a 25% undercount. This happens at two locations in the droplet, indicated by highlighted patches. At 270min a bacterium can also be seen to come close to the droplet boundary (shown by highlighted patch). The manual count differs by 1 depending on whether this is counted as being inside the droplet or not.

**TRACK B.**
**Orange rectangle**

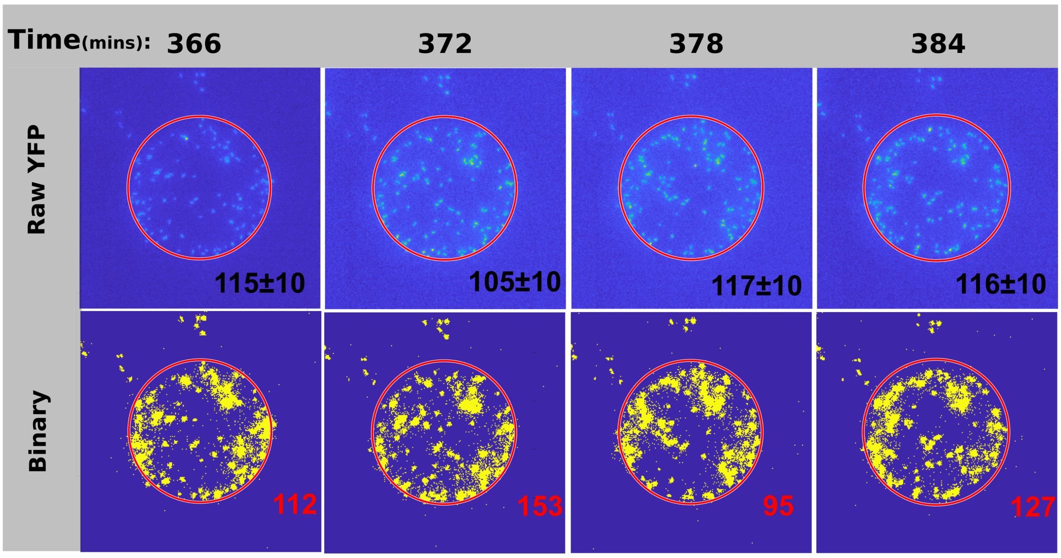


FIG. S6: **Transient dip: example B**. Microscopy images (fluorescence image and binary thresholded image) from the trajectory shown in orange in Figure S4. The images correspond to the large dip in bacterial count at a late stage in the trajectory (372min; shaded orange area in Figure S4). The numbers on the images show the bacterial counts (black numbers are manual counts; red numbers are automated counts). The red circle shows the droplet boundary as detected by our software. In these images the density of bacteria in the droplet is very high, making both manual and automated counts difficult. The automated algorithm undercounts the bacteria due to clustering. Data from these late points in the trajectories was not used in our analysis because of the difficulty in obtaining accurate counts.

**Track C.**
**Yellow rectangle**

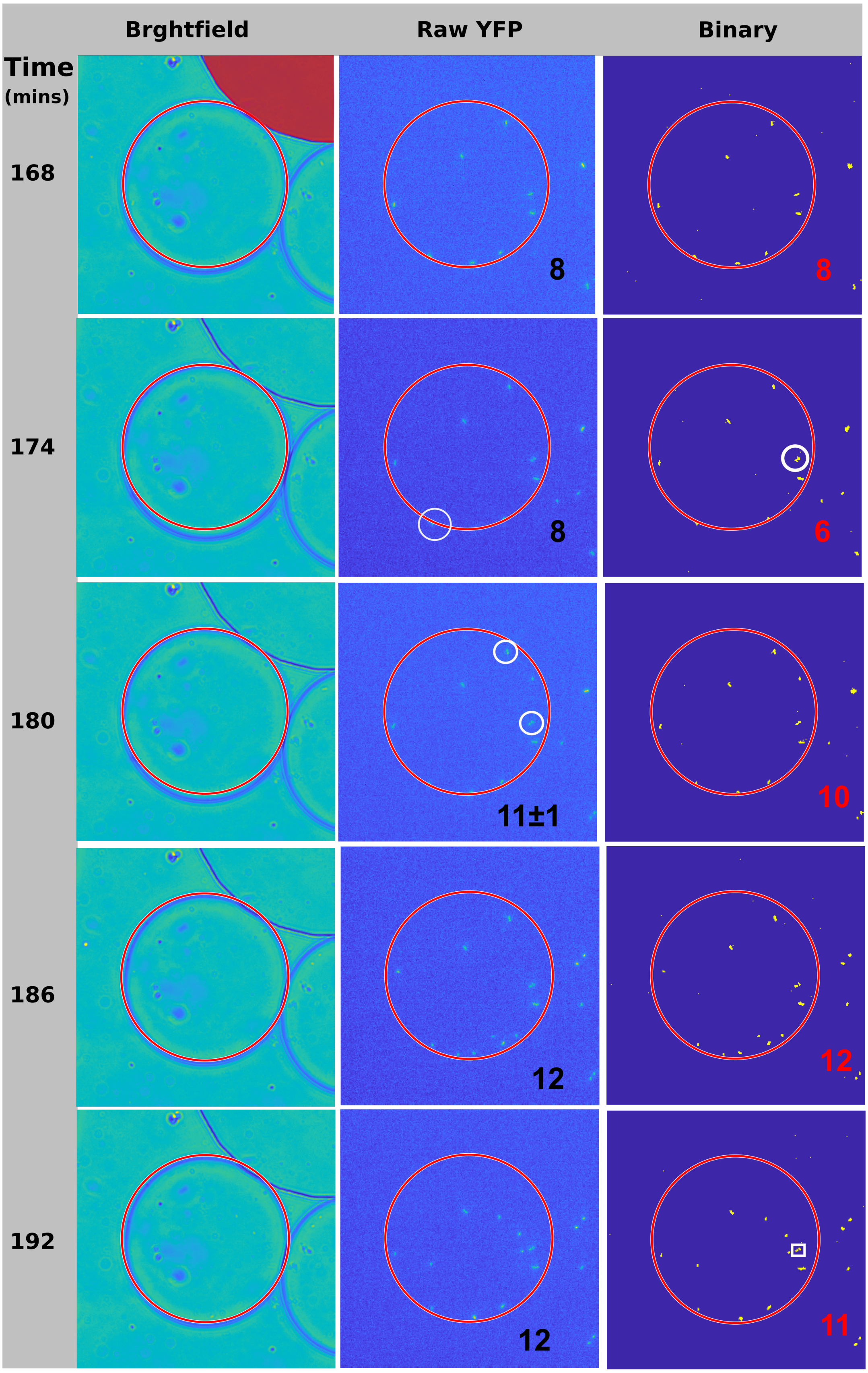


FIG. S7: **Transient dip: example C**. Microscopy images (fluorescence image and binary thresholded image) from the trajectory shown in yellow in Figure S4. The images correspond to the dip in bacterial count at 174min (shaded yellow area in Figure S4). The numbers on the images show the bacterial counts (black numbers are manual counts; red numbers are automated counts). The red circle shows the droplet boundary as detected by our software. This transient dip has multiple origins. Firstly, the droplet is somewhat distorted by the adjacent PDMS pillar (shaded red in the first time-frame) so that the boundary detection is inaccurate. This causes a bacterium to be missed in the count at 174min. Secondly, a pair of bacteria are co-localised in the same 174min image, and are counted as 1. Therefore the total count at 174min is 6 bacteria instead of 8. This colocalisation continues in the subsequent timestep (180min), and another pair of bacteria also co-localise, making it difficult to count even manually. Furthermore, a bacterium is missed in the image at 192min because the algorithm labels 8-connected objects, so the bacteria in the white square are counted as one.
